# Bio-inspired augmented reality: an interactive, digital twin of *C. elegans*

**DOI:** 10.1101/2024.05.29.596399

**Authors:** Daniel Sacristán, Sebastian Jenderny, Philipp Hövel, Christian Albers, Isabella Beyer, Karlheinz Ochs

## Abstract

This work presents a digital twin of the nematode *Caenorhabditis elegans* (*C. elegans*), an organism whose biology has been extensively studied. The digital twin can emulate neuronal activity and the corresponding muscle activity, and performs basic locomotion movement. The underlying mathematical model of *C. elegans* can be realized directly as an electronic circuit and is additionally implemented as a ready-to-use simulation in software. We implemented the digital twin in augmented reality (AR) as a novel format that extends the content of a traditional paper with an interactive visualization in the real world. The figures in the paper are the anchor point for the AR that can be accessed by the readers via an open-source app, which is freely available for tablets, phones, and AR glasses. This enables immersive experiences of the three-dimensional visualization in the real world from a perspective chosen by the reader, supplementing the traditional, flat figure layout of the paper. For researchers, the digital twin further provides a useful tool that is highly relevant and versatile for future developments. At the same time, its manifold possibilities for scientific outreach also aim at making the topic more engaging for a broader audience.

## 1 Introduction

Modern algorithms and artificial intelligence models can perform tasks such as pattern and language recognition with a similar or even higher accuracy than biological systems like the human brain [1, 2]. However, they require several orders of magnitude more energy than their biological counterparts to achieve similar performance [3, 4]. This is an obstacle for their widespread deployment and future development, since the amount of energy produced globally only grows at a rate of about 2% per year [5]. A very promising approach to tackle this problem are biologically-inspired artificial systems [3, 6].

Here, we present a digital twin of the nematode *Caenorhabditis elegans* (*C. elegans*), an organism that has been extensively studied [7–13]. The concept of a digital twin of *C. elegans* that can emulate locomotion behavior has also been explored in [14] by means of a very different approach based on using machine learning and an offline dataset to train a neural network consisting of 469 nodes and including 4869 chemical connections and 1433 electrical connections. Instead, we describe each neuron via the Hindmarsh-Rose model and propose a model for neuron behavior based on the equivalent electrical circuit from [15] and a locomotion model based on [16].

The proposed model of *C. elegans* can be realized directly as an electronic circuit [15] and is additionally implemented as a simulation in software. The advantages of simulating the electrical circuit with the wave digital concept are efficiency, modularity, numerical stability, real-time processing capabilities, potential for low computational complexity in certain applications, and ultimately a deeper understanding of the circuits as a didactic advantage. The digital twin emulates neuronal activity for 279 out of 282 neurons of the somatic nervous system of *C. elegans*. It also emulates the corresponding muscle activity for *C. elegans*’ 95 muscles, given sensory input, and is able to perform basic locomotion movement. In order to visualize the digital twin in an interactive manner, the simulation has been programmed in the Unity3D game engine [17]. That way, we can provide versions for computers, mobile devices, augmented reality (AR) [18–22], and virtual reality [23, 24]. The benefits of using immersive media for visualization of scientific data and as a tool for research have been widely researched [25–29]. The idea of using AR to display scientific data directly into publications has been studied in [30], but while the proposed web-based framework of that paper is well suited for online visualizations of data with reduced file-sizes, it is not suited for the complex interactive simulation of the digital twin proposed in the current study. This work allows to visualize the digital twin in a virtual and immersive environment, study the neuronal and muscular dynamics, and directly compare its behavior in real time to that of the biological counterpart. Thus, it provides a useful tool for both a general audience and researchers that is highly relevant and versatile for future developments.

## 2 Results

An overview of the presented digital twin concept and its working principles is depicted in Fig. 1. The digital twin is based on a mathematical model for the neuron, muscle, and locomotion activity that is translated into a software algorithm via methods of electrical engineering. Within the game engine Unity3D, this software algorithm feeds a visual model of the worm that builds upon the freely available 3D model of the Open Worm Project [31]. This 3D model is highly detailed and interactive as it allows to rotate, zoom in, and zoom out to explore the *C. elegans* from different angles. All implemented options are summarized in Tab. 1.

**Table 1:** Features (Real-time data and pre-recorded data modes).

| Name | Location | Description |
| --- | --- | --- |
| Rotate view | Main Window | Screen and dome: click & drag mouse<br>Handheld: drag screen with one finger<br>AR: move device |
| Zoom in/out | Main Window | Screen and dome: mouse-wheel up/down<br>Handheld: pinch in/out with two fingers<br>AR: move device closer/further |
| Export | Button, right-side menu | Export simulation results as csv file |
| Show/hide scale reference | Checkbox right-side menu | Show/hide the scale reference bar |
| Show all neurons | Checkbox right-side menu | Show all neurons with the same size |
| Show all connections | Checkbox right-side menu | Show/hide the whole connectome |
| Show no connections | Checkbox right-side menu | Disable the visualization of connections |
| Adjust skin transparency | Slider right-side menu | Set the transparency of the skin layer |
| Adjust neuron transparency | Slider right-side menu | Set the transparency of the neurons |
| Adjust muscle transparency | Slider right-side menu | Set the transparency of the muscles |
| Neuron Types | Legend on the bottom left | Color-coding of the neuron-groups |
| Close app | Button right-side menu | Exit the app |
| Tooltip information | Main Window | Click a neuron to select it and show the associated tooltip. Click again to deselect |

**Figure 1.**
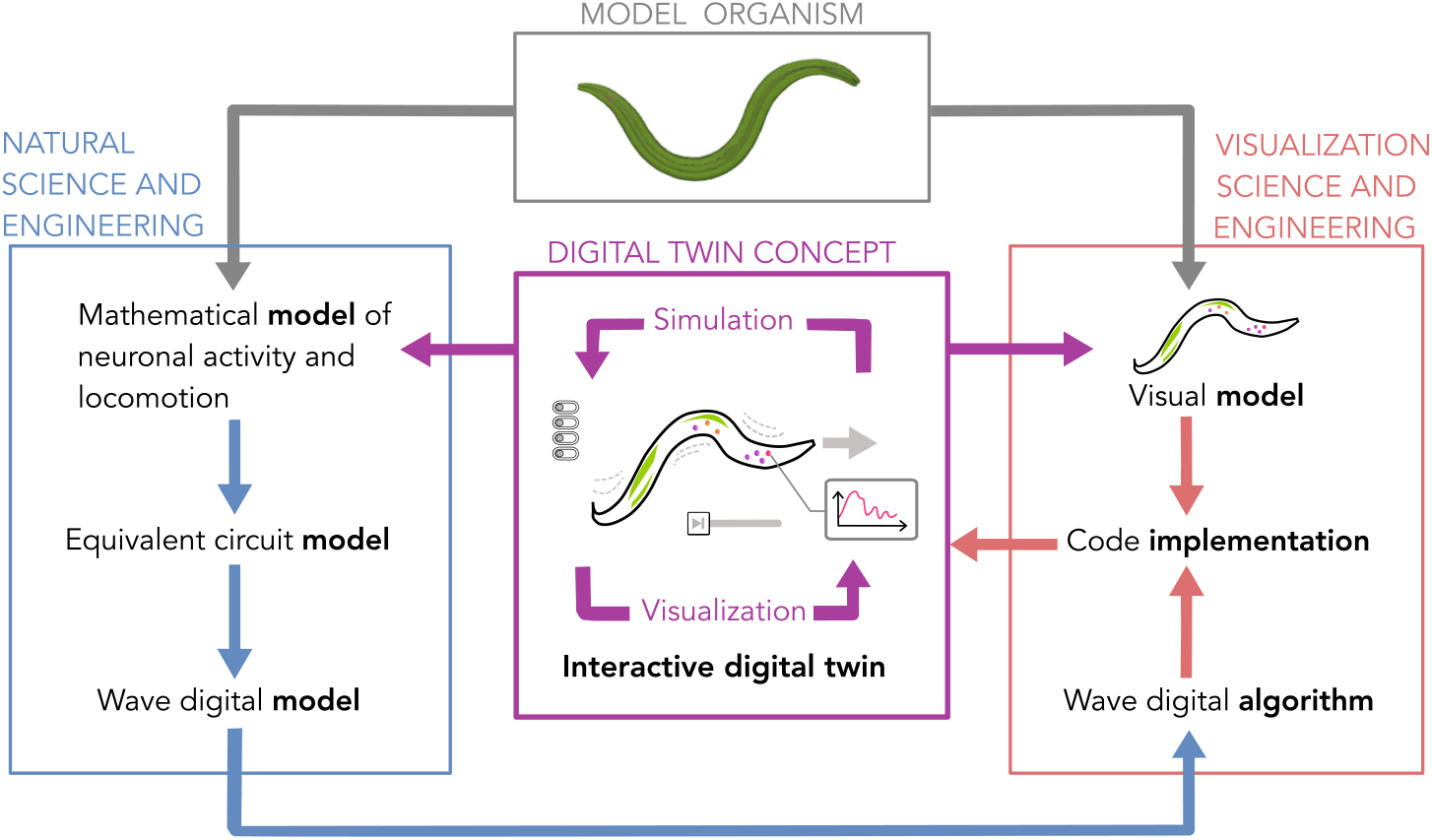
Digital twin concept, methodology and iterative working principle. As an interactive simulation and visualization platform at the interface of natural and visualization science and engineering, the digital twin can facilitate the development and research in the involved fields of knowledge.

Visualizations are made available for screens of computers or mobile devices, AR, and a 360^*°*^ dome view, see Tab. 2.

**Table 2:** Views.

| Name | Description | Purpose |
| --- | --- | --- |
| Screen | Desktop or handheld visualization | Traditional visualization on Windows and Android operating systems |
| AR | Augmented reality | Visualization of virtual objects in a real environment |
| Dome | 360 projection | For projection in domes and planetariums |

Results of the visualizations can either be obtained in real-time, with pre-recorded data, or with disabled visualization to only export simulation results, see Tab. 3. Details are provided in the following paragraphs.

**Table 3:** Modes of operation.

| Name | Description | Purpose |
| --- | --- | --- |
| Real-time data | The behavior of the digital twin is computed and displayed in real-time | Experiment with the digital twin |
| Pre-recorded data | The behavior of the digital twin is displayed using pre-recorded data | Demonstrate the system and run smoother on less powerful hardware |
| Computation only (Windows only) | Disabled Visualisation for faster computation | Fast export of the simulation results |

### 2.1 Visualization

Given an arbitrary input stimulus of the digital twin that leads to a forward locomotion, simulation results of the digital twin are visualized by an animation of the 3D model. Details of the mathematical and numerical model can be found in the supplementary material. To visualize the locomotion, each of the 24 transversal sections of *C. elegans* is animated according to its distance to the center axis of the digital twin. The result mimics the forward movement of the biological *C. elegans*. To better observe the different elements of the organism, the transparency of the skin and muscle layer can be adjusted. A screenshot of the overall visualization can be see in Fig. 2. One aspect of this visualization is the connectome of *C. elegans* within the 3D model. The neurons are color coded according to their functionality groups: interneurons (magenta), motor neurons (orange), sensory neurons (red), touch-anterior sensory neurons (blue), and touch-posterior neurons (cyan) [16]. Each individual neuron can be selected to highlight the neurons with which it is connected and the corresponding connections. Selecting a neuron also allows its name to be displayed in a tooltip together with a plot of the time series of its associated neural activity. This is depicted in Fig. 3(a).

**Figure 2.**
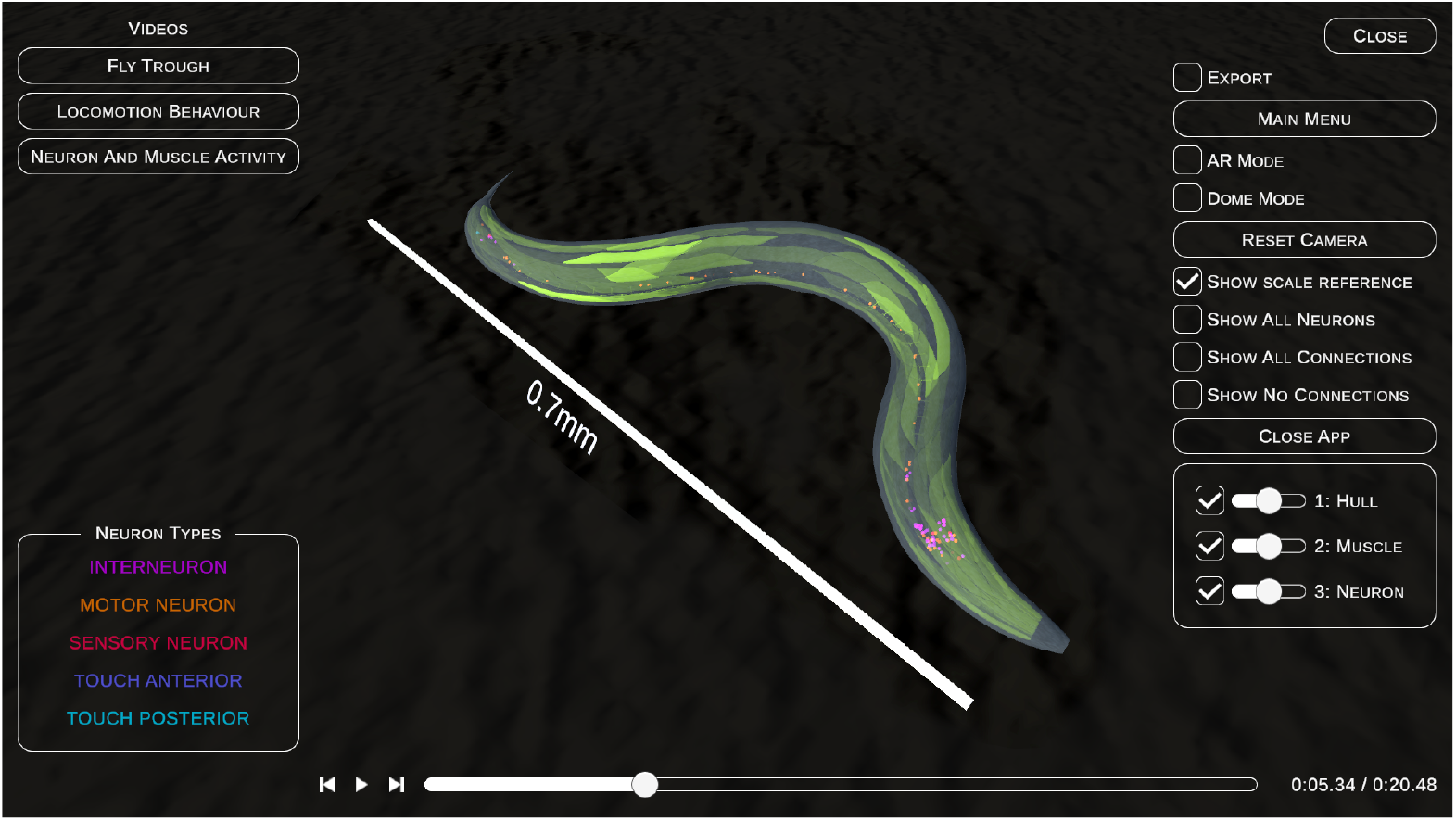
General view of the digital twin of *C. elegans*. The screenshot shows the graphical interface of the open-source app. The neurons are visualized by colored spheres as indicated by the legend, and the muscles are depicted by green color.

**Figure 3.**
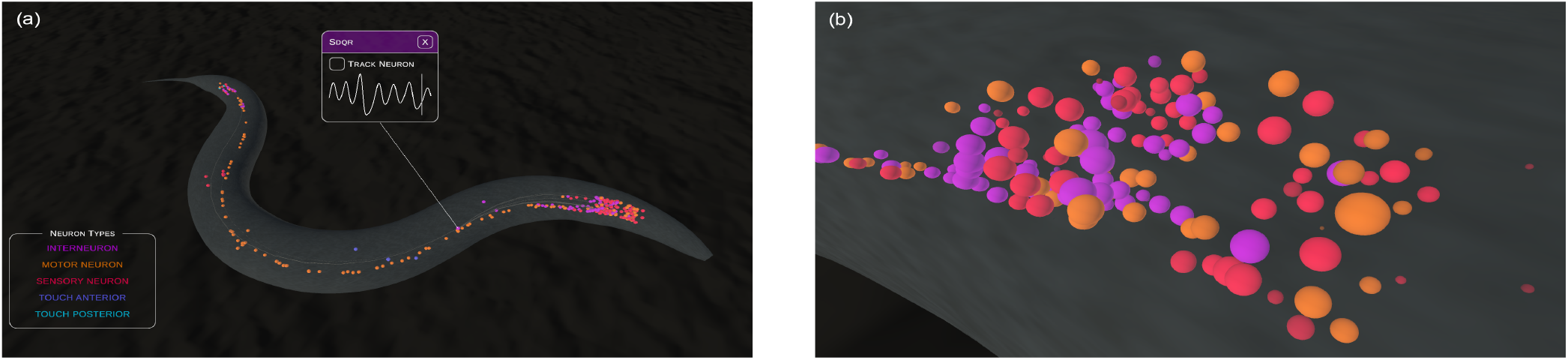
Connectome and neuron activity. (a) Visualization of the neural connectome in the digital twin of *C. elegans*. When a neuron is selected, its connections to other neurons are highlighted and a tooltip shows its name and a plot of its activity in time. (b) Visualization of the neuron activity in the digital twin of *C. elegans*. The size and brightness of each sphere represents the activity level of the corresponding neuron at a given time. The colors refer to the neuron type as seen in the legend depicted in the left panel.

Neuronal activity refers to the voltage of a neuron that is generated by the underlying mathematical model during a locomotion simulation. It is visualized through the size and brightness of the 3D representation of a neuron: A larger size and brighter color corresponds to a stronger neuronal activity, while a smaller size and darker color represents a weaker activity in the corresponding neuron (see Fig. 3(b)). Inactive neurons are represented as small spheres and with a darkest color. This is computed and represented in real time, and interaction with the neurons as described in the previous paragraph is also possible during the simulation.

Neuronal activity also drives muscle activity. In particular, each muscle is activated by those motor neurons to which it connects. Similar to the neuronal activity, muscle activity refers to the voltage that is calculated by the mathematical muscle model. In the 3D model, the muscle activity is visualized through the transparency of the muscle representation: Lower transparency corresponds to a stronger muscle activation and higher transparency represents a weaker activation of the corresponding muscle. Again, this is computed and represented in real time. An exemplary visualization of the muscle activity is shown in Fig. 4(a). A reference scale bar, as depicted in Fig. 4(b), can be shown or hidden and allows for a better comprehension of the scale of the organism in science dissemination activities.

**Figure 4.**
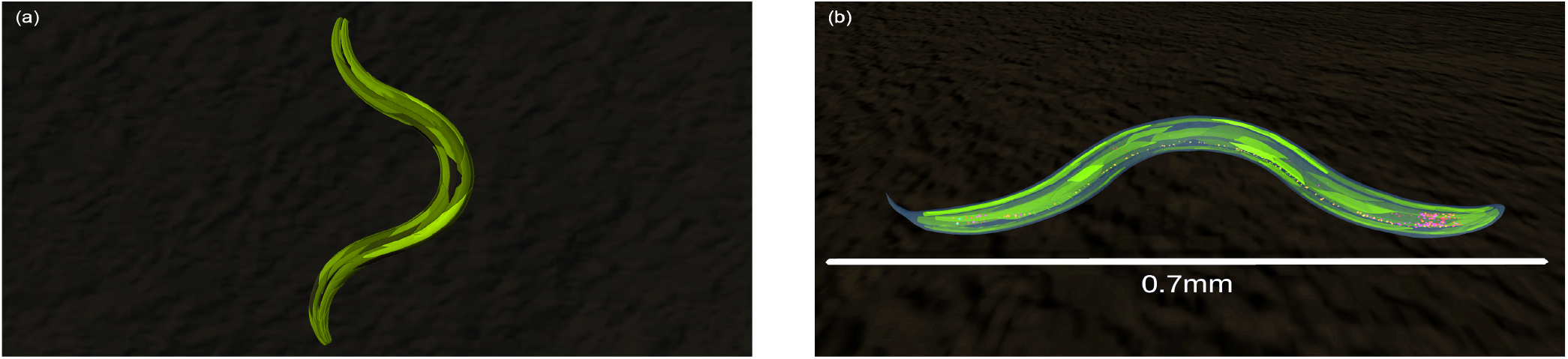
Muscle activity and reference scale bar. (a) Visualization of the muscle activity in the digital twin of *C. elegans*. The transparency level of each muscle represents its activity level at a given time instant. (b) Reference bar to illustrate the scale of *C. elegans*.

To cover for different needs when viewing the visualization of locomotion, connectome, as well as neuron and muscle activity, we have implemented user-driven interactions together with pre-set perspectives. Therefore, users can either move the camera and choose the view angle themselves, or select a pre-set perspective that shows the main features of the visualization in a curated way. The latter option, see Fig. 5, is particularly useful for dissemination purposes and scientific outreach, e.g., at expositions or presentations.

**Figure 5.**
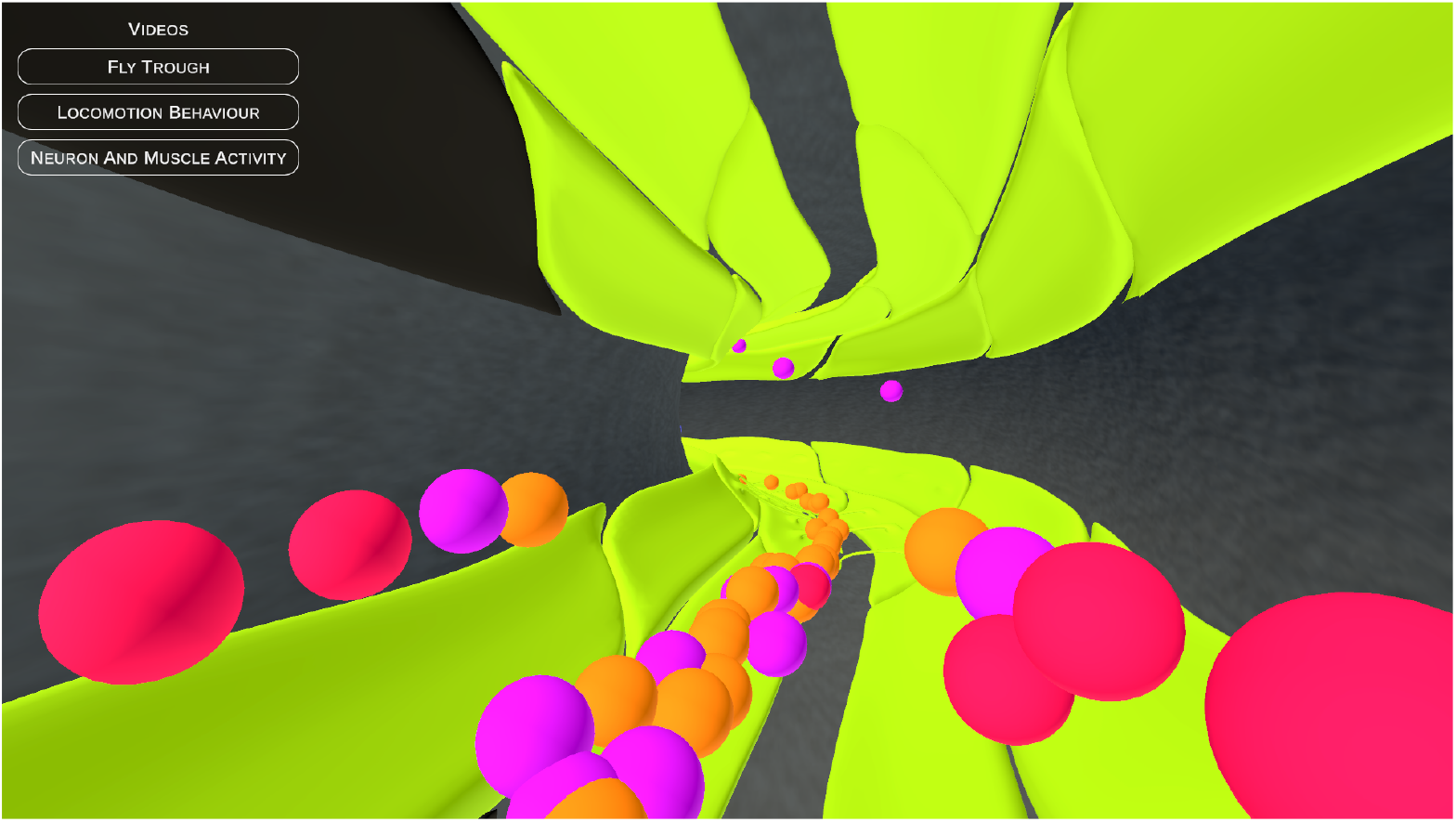
*Fly Through* perspective. Screenshot of the camera path inside the digital twin of *C. elegans*. The neurons are visualized by colored spheres (cf. Fig. 2), and the muscles are depicted by green color.

### 2.2 Supported View Formats

The digital twin is available for visualization on a computer screen (Windows operations systems), on Android devices, in Augmented Reality (AR), and in a 360 projection dome. A view of the AR mode is shown in Fig. 6(b) and is triggered by a QR code-like image as shown in Fig. 6(a). The app itself and its source code is available in an open access format. The dome view is illustrated in Fig. 6(c), while Fig. 6(d) shows the desktop screen view. The dome view is intended for large audiences in planetariums, while the AR view caters to museum exhibitions and mobile devices and the desktop mode targets smaller audiences and fixed traditional screens.

**Figure 6.**
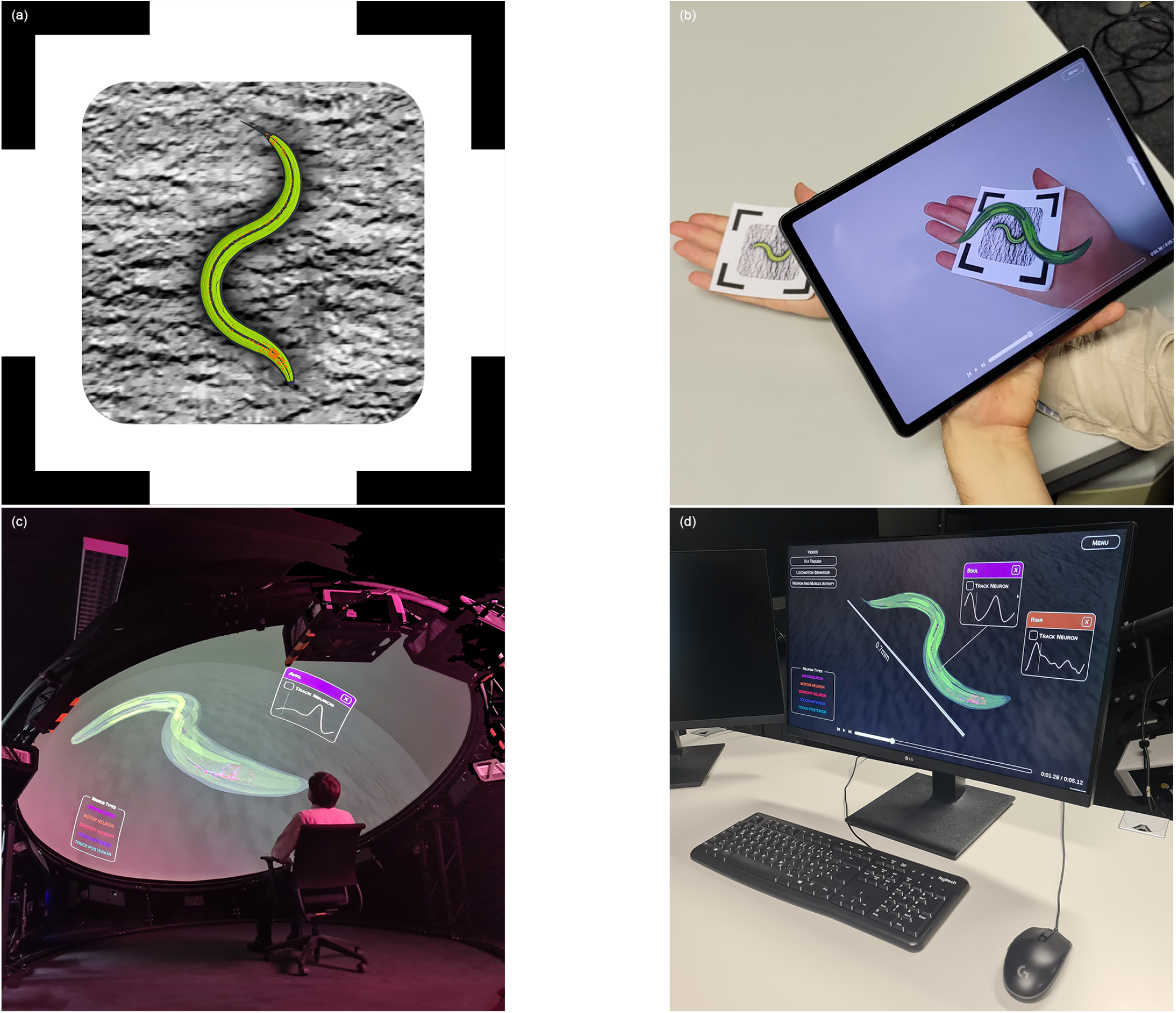
*Augmented Reality* mode, screen and dome views. (a) Image used to trigger and anchor the representation of the digital twin in augmented reality. (b) Augmented reality view. The 3D digital twin can be seen through the tablet. This allows the user to adjust the perspective by simply moving the device. (c) Dome view. (d) Screen view.

### 2.3 Modes of Operation

We have implemented three different modes of operation to cover for different purposes and audiences. The *real-time data* mode calculates both the mathematical model and the visualization in real-time. This especially allows for experiments with the digital twin, such as removing certain neurons or connections, whose effects can immediately be observed from the visualization.

Since our digital twin aims at general availability for a broad audience, we have implemented a second mode that uses only pre-recorded data. This makes the system run on low-performance hardware. In this *pre-recorded data* mode, the simulation is not computed in real-time. Instead, a previously recorded simulation is displayed, which lets the animation run smoothly on devices such as mobile phones and tablets, and is useful for demonstration and science dissemination activities.

As an additional feature, we implemented a *computation-only* mode that allows to use the software as a user-directed simulator. In this mode, the visualization is disabled to boost the computational performance. This allows the numerical integration of the model to run much faster since the computations for the visualization of the 3D model do not have to be performed. Upon completion of the simulation, the time series of all variables are available for export as comma-separated-value (csv) files and for subsequent analysis via the preferred methods of the user. This export feature is available for all three modes of operation.

## 3 Conclusions

The digital twin is a replica of the findings from biology and enables an immersive, experimental experience outside of the wet laboratory. The spectrum of applications ranges from a playful introduction to the topic for learners and the general audience to a promising tool for research by scientists. It can be used in research to, for instance, simulate and evaluate the effect of adding or removing individual neurons and/or connections. The digital twin facilitates experimental replication and enables systematic automatic exploration of modifications in the electrical or chemical layers of the connectome to identify the crucial parts for certain behavioral patterns. This way, the digital twin gives important insights that would be otherwise not or only with full-scale experimental equipment achievable.

Furthermore, the visualization of the digital twin allows researchers to observe the responses at a much more detailed level and from user-directed perspectives than *in vivo* experiments with its biological counterpart, which has practical limitations. For example, in the digital twin the point of view of the camera can be put inside the organism and objects such as organs or muscles can be hidden in the visualization in real time. Similarly, the performance of multiple neurons and muscles can be monitored simultaneously in a very convenient way.

Moreover, a *computation-only* mode enables direct simulations, including an export of the output data without representing it visually. This is particularly useful in iterative optimization approaches that have a numeric target and where a visualization of the locomotion animation is not needed for each iteration. This interactivity makes the digital twin particularly useful for researchers due to the possibility of running experiments systematically and without having the actual biological organism in the lab. Such *in silico* experiments can be easily replicated.

Finally, let us close with a brief outlook: The presented tool reads the information on the neurons’ relative positions and the different layers of connectome from external csv files. Therefore, the app can – in principle – be extended to other organisms as well. To invite other developers and researchers, we provide the source code in an open-access format under a BY-NC-SA creative commons licence.

## Methods

The methodology behind the presented digital twin is depicted in Fig. 1. At the heart is a mathematical model that describes neuronal activity, which drives muscle cells via a set of differential equations. These equations reflect an equivalent electrical circuit and can be translated into a wave digital model. For details, see Supplementary Information. The wave digital concept was originally developed to mimic electrical circuits with real-time algorithms on a digital processor. In order to achieve this, an electrical circuit is port-wise decomposed into sources and elements, where the latter can be one-ports or multiports. The remaining Kirchhoff interconnection network is also divided into elements like series and parallel connections. All these sources and elements have a direct correspondence to a wave digital counterpart, where linear combinations of voltages and currents appear as signals in the wave digital domain. Reactive elements as capacitors and inductors require a numerical integration oft the underlying differential equations. As it turns out, the trapezoidal rule as a best choice to preserve the energetic properties of these devices is commonly used.

## Data availability

The source-code for the app presented in this paper and other findings of this study are available from the corresponding author upon reasonable request.

## Acknowledgements

This work was funded by the Deutsche Forschungsgemeinschaft (DFG, German Research Foundation) - Project-ID 434434223 - SFB 1461.

We acknowledge Kerstin Meurisch for the co-development and creation of Fig.1.

## Author contributions statement

All authors designed the study. Daniel Sacristán, Christian Albers, and Isabella Beyer implemented the visualization. Sebastian Jenderny, Philipp Hövel, and Karlheinz Ochs developed the mathematical model. All authors discussed the results and wrote the manuscript.

