## Supplementary Information for "Bio-inspired augmented reality: an interactive, digital twin of *C. elegans*"

#### Appendix A Mathematical Model

The mathematical model that feeds the visualization describes neuronal activity that drives muscle cells. It serves as a basis for the implemented algorithm of the digital twin and is described in the following subsections.

##### A.1 Neuron Model

We describe each neuron $N \in [1, n]$ via the Hindmarsh-Rose model. A model for all coupled neurons then yields

$$\dot{\boldsymbol{z}}_1 = -\left[\alpha \boldsymbol{z}_1^2 - \beta \boldsymbol{z}_1\right] \boldsymbol{z}_1 + \boldsymbol{z}_2 - \boldsymbol{z}_3 + \boldsymbol{x} - G_{\mathrm{el}} \boldsymbol{\Lambda}_{\mathrm{el}} \boldsymbol{z}_1 \,, \quad \dot{\boldsymbol{z}}_2 = -\gamma \boldsymbol{z}_1 \circ \boldsymbol{z}_1 - \boldsymbol{z}_2 + c \,, \quad \frac{1}{\epsilon s} \dot{\boldsymbol{z}}_3 = \boldsymbol{z}_1 - \frac{1}{s} \boldsymbol{z}_3 - \phi_r \,, \tag{1}$$
$$\boldsymbol{x} = \boldsymbol{k} - G_{\mathrm{Glu}} \left[\boldsymbol{z}_1 - E_{\mathrm{exc}} \mathbb{1}\right] \circ \left[\boldsymbol{S}(\boldsymbol{z}_1) \boldsymbol{A}_{\mathrm{Glu}}\right] - G_{\mathrm{ACh}} \left[\boldsymbol{z}_1 - E_{\mathrm{exc}} \mathbb{1}\right] \circ \left[\boldsymbol{S}(\boldsymbol{z}_1) \boldsymbol{A}_{\mathrm{ACh}}\right]$$
$$- G_{\mathrm{GABA}} \left[\boldsymbol{z}_1 - E_{\mathrm{inh}} \mathbb{1}\right] \circ \left[\boldsymbol{S}(\boldsymbol{z}_1) \boldsymbol{A}_{\mathrm{GABA}}\right] \,, \quad \boldsymbol{S}(\boldsymbol{z}_1) = \left[\mathbf{1} + \mathbf{exp}\left(-\lambda \left[\mathbf{diag}(\boldsymbol{z}_1) - \theta \mathbf{1}\right]\right)\right]^{-1} \mathbb{1} \,,$$

with $\boldsymbol{z}_1, \boldsymbol{z}_2, \boldsymbol{z}_3, \boldsymbol{x}, \boldsymbol{k} \in \mathbb{R}^{n \times 1}$, $\boldsymbol{A}_{\mathrm{Glu}}, \boldsymbol{A}_{\mathrm{ACh}}, \boldsymbol{A}_{\mathrm{GABA}}, \boldsymbol{\Lambda}_{\mathrm{el}} \in \mathbb{R}^{n \times n}$. Here, $\boldsymbol{z}_1, \boldsymbol{z}_2$, and $\boldsymbol{z}_3$ denote the membrane potential, the recovery variable, and the bursting variable of the neurons, respectively. $\boldsymbol{x}$ is the input signal and is based on synaptic signals for the neurotransmitter types Glu, ACh, and GABA, as well as on a constant sensory signal $\boldsymbol{k}$. Here, all entries of $\boldsymbol{k}$ except for those that correspond to the touch sensory neurons PLML and PLMR are zero. Next, $\boldsymbol{S}(\cdot)$ is the synaptic activation function, $\circ$ is the Hadamard product, $\mathbf{1}$ is the unity matrix, $\mathbb{1}$ is a vector consisting of ones, and $\mathbf{diag}(\boldsymbol{z}_1)$ contains the elements of $\boldsymbol{z}_1$ as entries of a diagonal matrix. $\boldsymbol{A}_{\mathrm{Glu}}, \boldsymbol{A}_{\mathrm{ACh}}$, and $\boldsymbol{A}_{\mathrm{GABA}}$ are the adjacency matrices for the synaptic couplings of the corresponding neurotransmitter types, and $\boldsymbol{\Lambda}_{\mathrm{el}}$ is the Laplace matrix describing the gap junction connections. Parameters are chosen as follows: $\alpha = 10^6, \beta = 3 \cdot 10^3, c = 10^{-3}, \gamma = 5 \cdot 10^3, s = 4, \phi_r = -1.6 \cdot 10^{-3}, \epsilon = 0.005, G_{\mathrm{el}} = 0.2, G_{\mathrm{Glu}} = 0.1, G_{\mathrm{ACh}} = 0.1, G_{\mathrm{GABA}} = 0.15, E_{\mathrm{exc}} = 2 \cdot 10^{-3}, E_{\mathrm{inh}} = 2 \cdot 10^{-3}, \lambda = 10^3$, and $\theta = -0.25 \cdot 10^{-3}$.

##### A.2 Muscle Model

Following [1], we model each muscle $M \in [1, m]$ by a leaky integrator, which results in

$$\dot{\boldsymbol{z}}_4 = \frac{1}{\tau} \left[r \mathbb{1} - \boldsymbol{z}_4 + G_{\mathrm{ACh}} \boldsymbol{B}_{\mathrm{ACh}} \boldsymbol{z}_1 - G_{\mathrm{GABA}} \boldsymbol{B}_{\mathrm{GABA}} \boldsymbol{z}_1\right] \,, \quad \boldsymbol{z}_4 \,, \in \mathbb{R}^{m \times 1} \,, \quad \boldsymbol{B}_{\mathrm{ACh}} \,, \boldsymbol{B}_{\mathrm{GABA}} \in \mathbb{R}^{m \times n} \,, \tag{2}$$

with the muscle activity $\boldsymbol{z}_4$ and the coupling matrices $\boldsymbol{B}_{\text{ACh}}$ and $\boldsymbol{B}_{\text{GABA}}$ describing the neuro-musculular connections based on the corresponding neurotransmitter types. Parameters are set to $\tau = 100 \cdot 10^{-3}$ and $r = 10^{-3}$.

#### A.3 Generation of Locomotion via a Harmonic Wave Model

We determine the overall locomotion by calculating a harmonic wave for each dorsal and ventral muscle pair $P \in [1, p]$, and averaging the sum of all resulting harmonic waves, see [1]. The harmonic wave model reads

$$f(t) = \frac{1}{\max(f(P,t))} \sum_{P=1}^{p} f(P,t)\,, \qquad f(P,t) = \hat{f} \sin\left(\frac{2\pi}{l} - \omega(P,t_P)t + \phi(P,t_P)\right)\,, \tag{3}$$
$$\omega(P,t_P) = \frac{\pi}{t_P(k) - t_P(k-1)}\,, \qquad \phi(P,t_P) = \phi_0(P) + \sum_{k=1} \left[\omega(P,t_P(k+1)) - \omega(P,t_P(k))\right] t_P(k)\,,$$

with the amplitude $\hat{f} = 1$ and the wavelength $l = 18$. Note that the radian frequency $\omega$ and phase offset $\phi$ depend on time points $t_P$ at which extrema are found in the subtracted muscle activity $z_{4,\text{dorsal}} - z_{4,\text{ventral}}$. They are updated each time a new extremum is found for the corresponding muscle pair $P$.

### Appendix B Equivalent Electrical Circuit

Inspired by [2], we derive a vector-valued equivalent electrical circuit for the coupled neurons and muscles. This circuit is directly derived from the mathematical model and is used as a reference circuit for developing the digital model. The circuit is shown in Fig. 1 and is governed by

$$\begin{aligned}
C_0 \frac{\mathrm{d}\boldsymbol{u}_1}{\mathrm{d}t} &= -\boldsymbol{G}_1(\boldsymbol{u}_1)\boldsymbol{u}_1 + G_0\boldsymbol{u}_2 - \boldsymbol{i}_3 + \boldsymbol{j}_1 - \frac{I_0}{U_0} G_{\text{el}} \boldsymbol{\Lambda}_{\text{el}} \boldsymbol{u}_1\,, \quad \boldsymbol{G}_1(\boldsymbol{u}_1) = \left[\alpha \boldsymbol{u}_1 \circ \boldsymbol{u}_1 - \beta \boldsymbol{u}_1\right] G_0 \\
C_0 \frac{\mathrm{d}\boldsymbol{u}_2}{\mathrm{d}t} &= -G_0 \boldsymbol{u}_2 + \boldsymbol{j}_2 \\
L_3 \frac{\mathrm{d}\boldsymbol{i}_3}{\mathrm{d}t} &= \boldsymbol{u}_1 - R_3 \boldsymbol{i}_3 + \boldsymbol{e}_3\,, \quad L_3 = \frac{L_0}{\epsilon s}, \quad R_3 = \frac{1}{s G_0} \\
C_0 \frac{\mathrm{d}\boldsymbol{u}_4}{\mathrm{d}t} &= -G_0 \boldsymbol{u}_4 + \boldsymbol{j}_4(\boldsymbol{u}_1)\,, \quad \boldsymbol{u}_1 = \boldsymbol{z}_1 U_0\,, \quad \boldsymbol{u}_2 = \boldsymbol{z}_2 U_0\,, \quad \boldsymbol{i}_3 = \boldsymbol{z}_3 I_0\,, \quad \boldsymbol{u}_4 = \boldsymbol{z}_4 U_0\,, \\
\boldsymbol{j}_1 &= \boldsymbol{k} I_0 - G_{\text{Glu}} I_0 \left[\frac{\boldsymbol{u}_1}{U_0} - E_{\text{exc}} \mathbb{1}\right] \circ \left[\boldsymbol{S}\left(\frac{\boldsymbol{u}_1}{U_0}\right) \boldsymbol{A}_{\text{Glu}}\right] - G_{\text{ACh}} I_0 \left[\frac{\boldsymbol{u}_1}{U_0} - E_{\text{exc}} \mathbb{1}\right] \circ \left[\boldsymbol{S}\left(\frac{\boldsymbol{u}_1}{U_0}\right) \boldsymbol{A}_{\text{ACh}}\right] \\
&\quad - G_{\text{GABA}} I_0 \left[\frac{\boldsymbol{u}_1}{U_0} - E_{\text{inh}} \mathbb{1}\right] \circ \left[\boldsymbol{S}\left(\frac{\boldsymbol{u}_1}{U_0}\right) \boldsymbol{A}_{\text{GABA}}\right] \\
\boldsymbol{j}_2 &= -\gamma \frac{I_0}{U_0^2} \boldsymbol{u}_1 \circ \boldsymbol{u}_1\,, \quad \boldsymbol{e}_3 = -\phi_r U_0 \mathbb{1}\,, \quad \boldsymbol{j}_4 = I_0 \mathbb{1} + G_{\text{ACh}} \frac{I_0}{U_0} \boldsymbol{B}_{\text{ACh}} \boldsymbol{u}_1 - G_{\text{GABA}} \frac{I_0}{U_0} \boldsymbol{B}_{\text{GABA}} \boldsymbol{u}_1.
\end{aligned} \tag{4}$$

Here, $U_0 = 1\,\text{mV}, I_0 = 1\,\text{mA}, C_0 = 1\,\text{F}, L_0 = 1\,\text{H}$, and $G_0 = 1\,\text{S}$ are normalization constants.

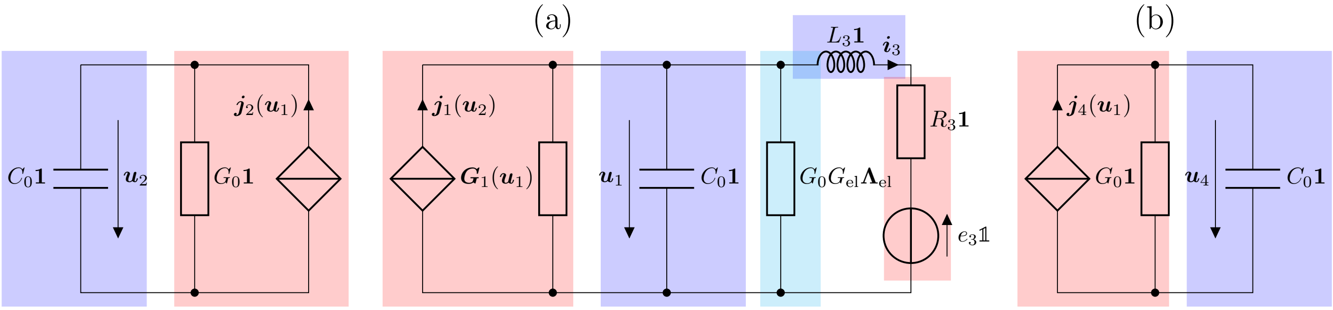


Figure 1: Vector-valued equivalent electrical circuit for (a) the coupled Hindmarsh-Rose models, and (b) the muscle models.

### Appendix C Wave Digital Model

Let us now translate the equivalent circuit from Fig. 1 into a wave digital model. Such a model constitutes a run-time efficient simulation algorithm mainly consisting of additions and multiplications. The translation of our equivalent circuit is done in two steps: (i) Decompose the circuit into its multi-ports, (ii) translate each multi-port as well as their interconnection structures into the wave digital domain using the bijective transformation

$$\boldsymbol{a} = \boldsymbol{u} + \boldsymbol{R}\boldsymbol{i}\,, \quad \boldsymbol{b} = \boldsymbol{u} - \boldsymbol{R}\boldsymbol{i}\,, \quad \boldsymbol{R} > \boldsymbol{0}\,, \tag{5}$$

see [3]. Here, $\boldsymbol{a}$, $\boldsymbol{b}$, and $\boldsymbol{R}$ are the incoming waves, the reflected waves, and the port resistances at a port of the equivalent circuit. Note that $\boldsymbol{R}$ is a diagonal matrix containing the scalar port resistances of the individual circuit elements of the vector-valued port.
Applying the transformation to the equivalent circuit, the capacitors and inductors translate to delay elements, and resistive sources translate to wave sources, given that the port resistances are chosen as shown in Eqn. 6:

$$\boldsymbol{R}_{C_1} = \frac{T}{2C_1}\boldsymbol{1}\,, \quad \boldsymbol{R}_{C_2} = \frac{T}{2C_2}\boldsymbol{1}\,, \quad \boldsymbol{R}_{C_m} = \frac{T}{2C_m}\boldsymbol{1}\,, \quad \boldsymbol{R}_{L_3} = \frac{2L_3}{T}\boldsymbol{1}\,, \quad \boldsymbol{R}_{j_2} = \boldsymbol{R}_{j_m} = \frac{1}{G_0}\boldsymbol{1}\,, \quad \boldsymbol{R}_{e_3} = R_3\boldsymbol{1}\,. \tag{6}$$

This leads to the reflected waves being expressed via

$$\begin{aligned}&\boldsymbol{b}_{C_\nu}(k) = \boldsymbol{a}_{C_\nu}(k-1)\,, \quad \boldsymbol{b}_{L_3}(k) = -\boldsymbol{a}_{L_3}(k-1)\,, \quad \boldsymbol{a}_{j_2} = \boldsymbol{R}_{j_2}\boldsymbol{j}_2(\boldsymbol{u}_1)\,, \quad \boldsymbol{b}_{j_m} = \boldsymbol{R}_{j_m}\boldsymbol{j}_m(\boldsymbol{u}_1)\,, \quad \boldsymbol{a}_{e_3} = e_3\mathbb{1}\,,\\ &\boldsymbol{a}_{j_1} = [\boldsymbol{1} + \boldsymbol{\rho}_j]\,\boldsymbol{j}_1(\boldsymbol{u}_2) + \boldsymbol{\rho}_j\boldsymbol{b}_{j_1}\,, \quad \boldsymbol{\rho}_{j_1} = [\boldsymbol{1} - \boldsymbol{G}_1(\boldsymbol{u}_1)\boldsymbol{R}_{j_1}]\,[\boldsymbol{1} + \boldsymbol{G}_1(\boldsymbol{u}_1)\boldsymbol{R}_{j_1}]^{-1}\,.\end{aligned} \tag{7}$$

The resistance $\frac{I_0}{U_0}G_{\mathrm{el}}\boldsymbol{\Lambda}_{\mathrm{el}}$ results in a multiplication with the scattering matrix

$$\boldsymbol{S} = [\boldsymbol{1} + \boldsymbol{\Lambda}_{\mathrm{el}}]^{-1}\,[\boldsymbol{1} - \boldsymbol{\Lambda}_{\mathrm{el}}]\,, \boldsymbol{R}_s = \frac{I_0}{U_0}G_{\mathrm{el}}\boldsymbol{1}\,, \tag{8}$$

where $\boldsymbol{R}_s$ is the associated port resistance. Finally, parallel and series interconnection translate to parallel and series adaptors that contain multiplier coefficients. The complete wave digital model is shown in Fig. 2 and can be implemented in software as follows: First, the values for the delay elements are initialized. Then, in an iterative manner, the incoming and the reflected waves of the parallel and series adaptors are computed. Note that we solve implicit relationships via additional fixed-point iterations [4]. For more details on wave digital modeling, the interested reader is referred to [3], [5].

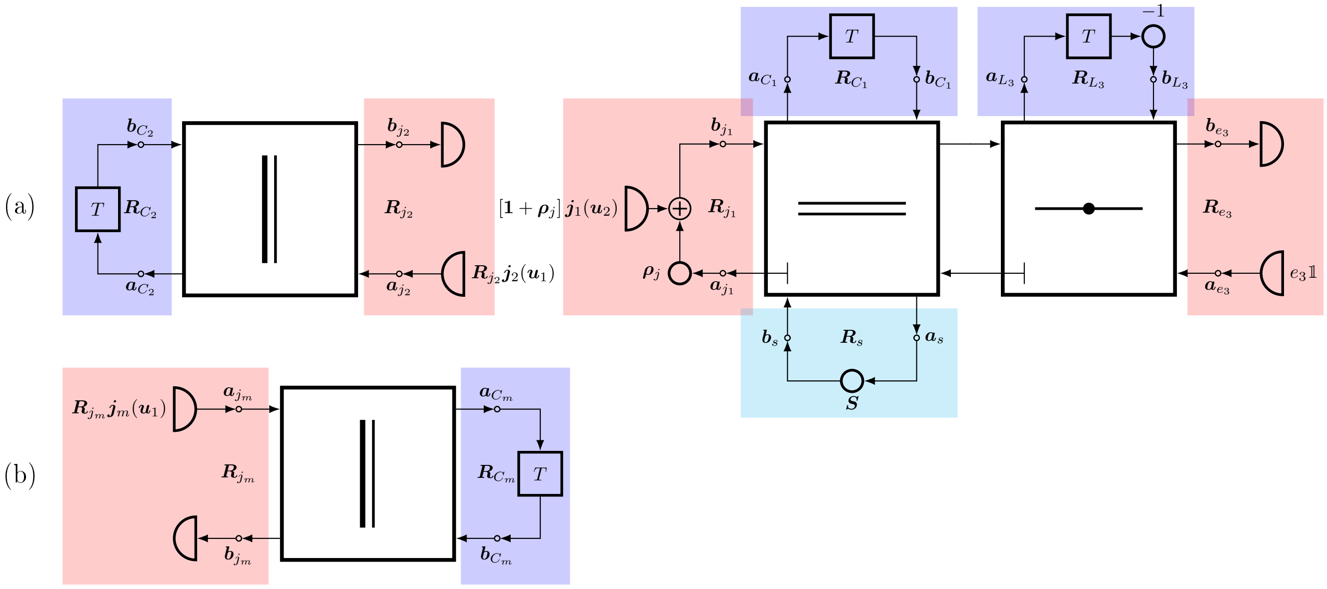


Figure 2: Vector-valued wave digital model for (a) the Hindmarsh-Rose circuit and (b) the muscle circuit.
